# Pseudoperplexity Probes Memorization in Protein Language Models

**DOI:** 10.64898/2026.06.08.730685

**Authors:** Alexander Plaikner, Manuel Ploner, Zeno Sewald, Tobias Senoner, Sebastian Franz, Maurice Brenner, Michael Heinzinger, Burkhard Rost

## Abstract

Protein Language Models (pLMs) have significantly advanced computational biology. Yet their scale and reliance on redundant training data raise a fundamental question: do pLMs generalize the statistical *grammar* of proteins, or do they simply memorize their training data? To investigate this, we used pseudoperplexity as a probe for sequence-level memorization, comparing ProtT5′s pseudoperplexity on a pre-training proxy dataset against a post-training holdout of genuinely novel sequences. To ensure a valid comparison, we matched the datasets by sequence length, cluster size, and taxonomic family. As a statistical baseline, we trained *n*-gram language models; analysis of higher-order *n*-gram composition and a statistically significant divergence in perplexity confirmed that the post-training sequences were genuinely novel at the local sequence level. ProtT5 showed a statistically significant difference in pseudoperplexity between seen and unseen sequences, though further analysis revealed this memorization signal to be modest. These findings suggest that ProtT5 exhibits detectable but limited memorization of its training data as measured by a pseudoperplexity-based probe.

## 1. Introduction

In recent years, protein Language Models (pLMs) have fundamentally transformed computational biology and downstream experimental applications relying on protein sequences, structures, and function^1–1^. By applying the transformer architecture^1^, which proved powerful in natural language processing (NLP) to protein sequences, models such as ProtT5^1^ and ESM-^1^ have learned rich representations of protein sequences from large corpora of unlabeled sequences. ProtT5-XL-U50, an encoder-decoder model with approximately three billion parameters based on the T5 architecture^1^, was trained on UniRef50^1^ and the Big Fantastic Database (BFD)^1,1^ with over 300 billion amino acids. Leveraging these representations, pLMs have enabled significant advances in protein prediction without having to rely on costly multiple sequence alignments (MSAs).

The scale of these pLMs raises the fundamental question: to what degree do they generalize by understanding the language of life^1^, i.e., the *grammar* of protein sequences, and to what extent do they simply memorize their training data. In the NLP domain, large language models (LLMs) can memorize substantial portions of their training data^15,16^, with memorization scaling with model capacity and data duplication^17^. Deduplication of training data has been shown to reduce memorization substantially^18^. Importantly, larger models tend to memorize data more readily, and masked language models (MLMs) overfit faster than autoregressive models when training data is limited or repeated^19^. This is directly relevant for pLMs trained on MLM objectives using clustered protein databases with inherently constrained sequence diversity^19^.

The distinction between memorization and generalization has direct practical consequences. Hermann et al. demonstrated that data leakage from pLM pretraining can distort downstream benchmark performance by approximately 11%^20^. A pLM that has memorized its training data would incorrectly assign high confidence to similar sequences while producing inflated uncertainty estimates for novel proteins. This would compromise its reliability in tasks such as scoring the fitness of newly discovered proteins or predicting variant effects of existing ones. Moreover, as protein databases — particularly those derived from metagenomic data — grow rapidly, understanding how pLMs respond to previously unseen protein sequences is crucial for their continued utility.

Pseudoperplexity, first introduced by Salazar et al.^21^ as a score based on the pseudo-log-likelihood (PLL) for bidirectional masked language models, provides a way to assess a model’s uncertainty about an input sequence. Low pseudoperplexity indicates that the model assigns high pseudo-log-likelihood to the observed sequence, indicating that the sequence conforms to the statistical patterns learned during training. Recent research demonstrates that sequence likelihood, which quantifies the implicit preference a model learns during pretraining, is directly predictive of its zero-shot fitness estimation capabilities^22^. In practice, mutation effect prediction suffers when the wild-type sequences are either under-preferred or over-preferred by the model. Beyond its use for zero-shot fitness estimation^23,24^ and as a predictor of ESMFold^3^ output quality, pseudoperplexity can serve as a diagnostic for overtraining: a model that has memorized its training data will assign systematically lower pseudoperplexity to proteins similar to the training data than to those absent from the training distribution, i.e., those deposited after training. Conversely, the absence of such a difference would suggest that the model has learned to generalize the underlying statistical grammar of protein sequences, assigning comparable pseudo-log-likelihood to both training and novel proteins.

Here, we applied this reasoning to ProtT5 by constructing a temporal holdout experiment. We compared pseudoperplexity values between a pre-training proxy dataset of UniRef50 sequences that were already present before the ProtT5 training, and a post-training holdout of proteins that were introduced in the 2025_01 UniProt release but absent from the 2024_01 release, meaning ProtT5 could not have encountered them during training. To ensure a valid comparison, we also excluded sequences similar to the ones in the training set, and matched pre- and post-training datasets in terms of sequence length, cluster size, and taxonomic family.

To provide a statistical baseline that operates purely on local sequence statistics without any learned representations, we additionally trained *n*-gram language models for orders *n* = 1, …, 7 and evaluated their perplexity on both dataset splits. Complementary analysis with these *n*-gram models confirmed that the two datasets differ significantly in their higher-order *n*-gram distributions, supporting the conclusion that the post-training sequences are genuinely novel at the local sequence level.

Building on this verified holdout, we used pseudoperplexity as a probe for memorization and found a statistically significant difference between pre- and post-training datasets, indicating that ProtT5 assigns a systematically higher likelihood to seen sequences. An area under the receiver operating characteristic curve (AUC-ROC) analysis further quantified this memorization signal as modest. Together, these results provide strong evidence of detectable sequence-level memorization in ProtT5 under a pseudoperplexity-based probe.

## 2. Background

### 2.1. Perplexity and pseudoperplexity

Perplexity is a standard metric for evaluating the quality of a language model. For an autoregressive model that assigns probability *P* (*x*_*i*_| *x*_1_, …, *x*_*i*−1_) to each token given all preceding tokens, the perplexity of a sequence of length *N* is defined as the exponentiated average negative log-likelihood per token:

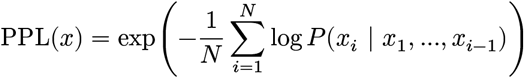

Perplexity can be interpreted intuitively as the average number of different tokens the model is choosing from at each position. A perplexity value equal to the vocabulary size *V* corresponds to a model that only randomly guesses the correct token, while lower values indicate greater certainty about the correct token. The smallest possible perplexity value is 1, corresponding to a model that assigns probability 1 to the correct token at every position.

Masked language models such as ProtT5 are not autoregressive and therefore the above perplexity calculation cannot be used directly. Salazar et al.^1^ introduced the pseudo-log-likelihood (PLL) score as an adaptation of perplexity to bidirectional masked language models. The pseudoperplexity of a sequence is computed by masking each position *i* in turn while keeping all other positions visible, querying the model for the probability it assigns to the correct residue (amino acid in protein referred to as ‘residue’) at that position, and accumulating these log probabilities:

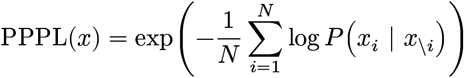

Here *x*_\*i*_ denotes the full sequence with position *i* masked. Because the model has access to both left and right context at every position, pseudoperplexity benefits from bidirectional information and will therefore be systematically lower than autoregressive perplexity for the same sequence. Low pseudoperplexity indicates that the model assigns high probability to the correct residue in a sequence. This translates to the sequence being familiar and is consistent with the statistics the model has learned during training.

### 2.2. N-gram language models

*N*-gram language models are among the simplest probabilistic models usable on sequences. An *n*-gram model estimates the probability of a token *x*_*i*_from the *n* − 1 immediately preceding tokens, approximating the full sequence probability as a product of these local conditional probabilities^1^. The conditional probability of a token *x*_*i*_in an *n*-gram is estimated from dataset counts as:

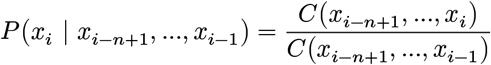

Here *C*(⋅) denotes the count of a given *n*-gram in the training dataset. A fundamental challenge for higher-order *n*-gram models is data sparsity: as *n* increases, many possible *n*-grams are never observed in the training dataset, leading to zero counts if previously unobserved *n*-grams appear in a validation dataset. Laplace (add-one) smoothing addresses this by adding a pseudocount of 1 to every *n*-gram^1^. This ensures that all counts are > 0 and that all sequences receive a non-zero probability even if they were never seen during training:

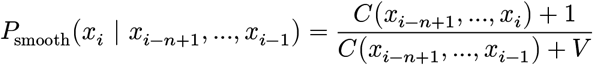

Here *V* is the vocabulary size. Despite their simplicity, *n*-gram models provide an interpretable reference point. Their perplexity reflects only local compositional statistics, since they lack the capacity to capture long-range dependencies like attention-based transformer models.

## 3. Methods

### 3.1. Dataset preparation

To evaluate whether a memorization signal is detectable in ProtT5, we constructed two dataset splits: a pre-training proxy, representing sequences the model was exposed to during training, and a post-training holdout, comprising sequences deposited in the database after the model’s training cutoff, redundancy-reduced against the training data, and therefore absent from it.

#### 3.1.1. Pre-training dataset

We derived the pre-training dataset from the UniRef50 release 2019_0112, which ProtT5 used during training and contains 28,919,163 cluster representatives in total. We then joined these sequences against the UniRef50 2026_01 release, from which we sourced all metadata: we retained only sequences that are cluster representatives in the 2026_01 release, recording their 2026_01 cluster size, PE (Protein existence) level, sequence uncertainty annotations, and taxonomic family, where the latter was obtained using `taxopy`^1^ together with the NCBI taxonomy^1^ dump from 2026-04-01.

We retained only sequences satisfying PE levels 1 or 2, where PE level 1 requires experimental evidence at protein level and PE level 2 requires experimental evidence at transcript level. The only filtering rule applied to the raw amino acid sequences from the 2019_01 release was the removal of sequences containing non-standard residue codes (B, J, O, U, Z, X). Together with the removal of sequences flagged with sequence uncertainty annotations in the 2026_01 release, this ensures that no uncertain or non-standard residues are present in the dataset, as retaining such residues would conflate model uncertainty with sequence-level ambiguity, making it impossible to attribute any observed pseudoperplexity signal purely to model behavior. After these filtering steps, 205,072 sequences remained.

#### 3.1.2. Post-training dataset

The post-training dataset served as the temporal holdout. It contains 1,237 proteins that were introduced in the UniProt28 release 2025_01 but were absent from the 2024_01 release^1^, meaning ProtT5 could not have encountered them during training. Following the same procedure as for the pre-training dataset, we joined these sequences against the UniRef50 2026_01 release to source all metadata, recording their cluster size, PE level, sequence uncertainty annotations, and taxonomic family. From the 2026_01 metadata, we retained only sequences that are cluster representatives in the 2026_01 release, satisfy PE levels 1 or 2, and are not flagged with sequence uncertainty. We additionally inspected the raw amino acid sequences from the 2025_01 release and removed those containing non-standard residue codes (B, J, O, U, Z, X).

As mentioned previously, ProtT5 was first trained on the BFD and subsequently fine-tuned on UniRef^1^. To ensure that the post-training dataset does not contain any data similar to either of these training sources, we removed all post-training sequences with more than 30% sequence identity and 90% coverage to any entry in the reduced BFD database^1^, which consists of the first non-consensus sequence from each BFD alignment, as well as to any entry in the UniRef50 2019_01 release. Both filtering steps were performed using MMseqs2^1^. After these filtering steps, 358 sequences remained.

#### 3.1.3. Sequence length and cluster size filtering

To reduce computational overhead and ensure compatibility, we applied the following filtering criteria to both datasets individually:

1. **Sequence length**: Because pseudoperplexity computation scales cubically with sequence length, we constrained the length of proteins to 1,000 residues.
2. **Cluster size**: The distribution matching step described in Section 3.1.5 uses stratification bins with a maximum cluster size of 100. Pre-training sequences with a cluster size above 100 would therefore never be selected as a match and were removed to reduce the size of the pre-training dataset before matching.

After filtering, 166,116 sequences remained in the pre-training dataset and 308 in the post-training dataset.

#### 3.1.4. Family filtering

Due to a large class imbalance in taxonomic family representation, we pruned overrepresented families in the post-training dataset. Several large families, such as Halisarcidae, Tarsonemidae, Limacodidae, and Elaeagnaceae, were almost exclusively present in the post-training dataset, making it impossible to find suitable matches in the pre-training dataset. For this reason, we removed all families from the post-training dataset that had no matching counterpart in the pre-training dataset. For families that were overrepresented in the post-training dataset but had at least one matching entry in the pre-training dataset, we pruned the post-training family size to at most twice the number of pre-training entries for that family. This was done to retain a larger number of sequences, as exact matching would have yielded 5.8% fewer sequences. Among the pruned entries, we retained those sequences that best matched the pre-training sequences of the same family in terms of sequence length and cluster size, to facilitate the distribution matching described in the next step. For example: if family A contained 72 entries in the post-training dataset but only 5 in the pre-training dataset (2 * 5 ≪ 72), we pruned the post-training family size to 2 × 5 = 10, retaining the best sequences as described previously. After filtering, 166,116 sequences remained in the pre-training dataset and 191 in the post-training dataset.

#### 3.1.5. Distribution matching

To eliminate possible confounders, we matched the pre-training dataset to the post-training dataset along three strata:

1. **Taxonomic family**

2. **Sequence length**: [0, 50], (50, 100], (100, 150], (150, 200], (200, 300], (300, 400], (400, 500], (500, 600], (600, 800], (800, 1000]

3. **Cluster size**: (0, 1], (1, 5], (5, 10], (10, 20], (20, 50], (50, 100]

The choice of bin sizes was motivated by the underlying density of sequence lengths and cluster sizes in the unfiltered datasets, as shown in Figure S2 and Figure S3.

For each post-training sequence, we selected a unique pre-training counterpart without replacement. When no exact match was available, we relaxed the constraints incrementally following a hierarchy of eight rules, ordered from most to least restrictive:

1. Match: taxonomic family, sequence length, cluster size.
2. Match: taxonomic family, cluster size.
3. Match: taxonomic family, sequence length.
4. Match: taxonomic family.
5. Match: sequence length, cluster size.
6. Match: cluster size.
7. Match: sequence length.
8. Match: nothing (uniform random selection).

The hierarchy reflects the strictness of each stratum: taxonomic family is placed first because it allows only exact matches, whereas sequence length and cluster size are defined over bins, meaning that any value falling within the same bin is considered an equally valid match.

The final distribution-matched pre-training and post-training datasets are equal in size, each containing exactly 191 sequences.

#### 3.1.6. Matched distributions

Figure S1a, Figure S1b, and Figure S1c show the result of the matching process for both datasets in terms of sequence length, cluster size, and taxonomic family. Overall, the matching procedure was successful. The cluster size distribution aligns most precisely, with only a single difference between the matched-pre-training and post-training distribution. In total, across both datasets of 191 sequences each, 18 sequences could not be placed in the correct sequence length bin, 1 in the cluster size bin, and 13 in the correct taxonomic family. The limiting factor for taxonomic family matching was the class imbalance which was not fully resolved in Section 3.1.4 due to limited data. Families outside the top 10 most represented post-training families may have up to a factor of two imbalance, though since these families contain at most 4 sequences in the post-training dataset, the absolute difference in any such case is at most 2. The main limiting factor throughout the entire matching process was the taxonomic family stratum, as it allows only exact matches.

As an additional check for possible remaining confounders, we visualized the ProtT5 embeddings of the matched datasets with t-SNE and colored the same projection by sequence length, cluster size, and taxonomic family (Figure S5a, Figure S5b, and Figure S5c). Sequence length was partly reflected in the embedding space, with longer sequences tending to concentrate in the central region. However, because sequence length was explicitly included in the matching procedure, this length-related structure was expected to affect both matched datasets similarly. Cluster size and taxonomic family did not form clear spatial patterns.

### 3.2. N-gram baseline models

To complement our analysis of ProtT5 pseudoperplexity on the pre- and post-training datasets, we constructed an interpretable statistical baseline using *n*-gram language models, ranging from unigrams to 7-grams. By analyzing perplexity values derived from these simple count-based methods, we quantified differences in *n*-gram distributions between the pre- and post-training datasets and showed that these simple models begin to overfit as the context window size increases.

#### 3.2.1. Training dataset

All *n*-gram models were trained on the dataset described in Section 3.1.1. The *n*-gram training dataset contains 205,072 protein sequences totaling approximately 68 million tokens. It is important to note that this training dataset is a superset of the pre-training dataset we use when evaluating the perplexity values of the *n*-gram models. This intentional overlap mirrors our evaluation of ProtT5, allowing us to compare model behavior on a subset of its training data against previously unseen post-training sequences.

#### 3.2.2. N-gram construction and counts

To correctly construct *n*-grams of various orders, all sequences were padded prior to counting. Each sequence was prepended with *n* − 1 start-of-sequence tokens <s> and appended with a single end-of-sequence token </s>. The start tokens are required such that the conditional probability of residues near the beginning of a sequence can be computed correctly - for example, computing the probability of the first residue under a trigram requires a context of two <s> tokens. Since the <s> token is only used as context for other tokens, and therefore never generated, it is excluded from the vocabulary. The final vocabulary used for the *n*-gram models has a size of *V* = 21, including the 20 standard amino acids and the end-of-sequence token </s>.

*N*-gram counts are accumulated using a sliding window of width *n* over all padded sequences in the training dataset, separately for each order of *n*-gram. These counts fully define each model and are sufficient to calculate the probabilities required for perplexity computation.

#### 3.2.3. Probability estimation and perplexity calculation

The conditional probability of a token *x*_*i*_given its preceding context was calculated as the relative frequency of the full *n*-gram over all completions of that context as described in Section 2.2. To solve the issue of zero counts for unseen *n*-grams, we applied Laplace (add-one) smoothing for all models as described in Section 2.2.

The perplexity of a sequence, including the end-of-sequence token </s>, was computed as described in Section 2.1. This yields a measure of the model’s uncertainty over the full generative process, including sequence termination.

### 3.3. Bootstrap estimates

The comparison of perplexity and pseudoperplexity values between the pre- and post-training datasets was performed using a non-parametric bootstrap procedure, which we applied consistently across both the *n*-gram models and ProtT5.

For each *n*-gram order (*n* = 1, …, 7) as well as for ProtT5, and for each dataset, we drew 10,000 bootstrap resamples with replacement from the per-sequence perplexity or pseudoperplexity values and computed the mean for each resample, yielding an empirical distribution of bootstrap means.

To test for statistically significant differences between the two datasets at a given *n*-gram order, we computed the paired difference of bootstrap means (*µ*_pre_− *µ*_post_) across all 10,000 resamples. Because we performed seven such comparisons (one for each *n*-gram order from 1 to 7), we applied a Bonferroni correction to control the family-wise error rate at *α* = 0.05 (which coincides with a 95% confidence interval). A difference was declared statistically significant if the resulting 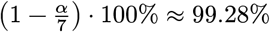 confidence interval for the difference did not contain zero.

Similarly, to evaluate whether increasing the *n*-gram order by one produced a statistically significant difference in perplexity within the same dataset, we computed the paired difference between the bootstrap means of consecutive *n*-gram orders (*µ*_*n*_− *µ*_*n*+1_). For each dataset, this involved six comparisons (orders 1-2, 2-3, …, 6-7). A Bonferroni correction with 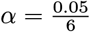 was therefore applied separately to each dataset, and the difference in perplexity was declared statistically significant if the corresponding 99.17% confidence interval did not contain zero.

For the ProtT5 pseudoperplexity analysis, we performed only a single comparison between pre-training and post-training datasets. A difference in perplexity was declared statistically significant if the 95% confidence interval of the bootstrap mean differences (*µ*_pre_− *µ*_post_) did not contain zero.

### 3.4. Memorization quantification

To quantify the extent of memorization, we used pseudoperplexity as a binary classifier to separate pre-from post-training sequences via a receiver operating characteristic (ROC) analysis. A model that has memorized parts of its training data is expected to assign lower uncertainty to familiar sequences than to novel ones. We exploit this property by sliding a threshold across the full range of pseudoperplexity scores, classifying all sequences below the threshold as pre-training and all sequences above as post-training. At each threshold value, we calculate the true positive rate (the fraction of pre-training sequences correctly classified as pre-training) and the false positive rate (the fraction of post-training sequences incorrectly classified as pre-training). Plotting the true positive rate against the false positive rate across all thresholds yields the ROC curve. The area under this curve (AUC) summarizes the classifier’s discriminative ability into a single scalar value, where 0.5 corresponds to random separation, indicating no detectable memorization, while a value of 1.0 would indicate perfect separation. We computed bootstrap confidence intervals for the AUC using the same 10,000-resample procedure described above, drawing sequences with replacement to create resampled datasets of the original size and computing the AUC for each resample.

## 4. Results and discussion

### 4.1. ProtT5 showed modest memorization

The pseudoperplexity distributions differed significantly between the pre- and post-training datasets (Figure 1a: pre-training: 9.07 with CI95 - 95% confidence interval - of (8.37, 9.79) vs. post-training: 10.18 with CI95 of (9.62, 10.75)). This suggested that ProtT5 has memorized some of its training data. The upper mode of both distributions was predominantly driven by short (< 50 residues) proteins (Figure S4). Possibly, short proteins share some property leading to higher pseudoperplexity, which may be just correlated or caused by their shortness.

**Figure 1:**
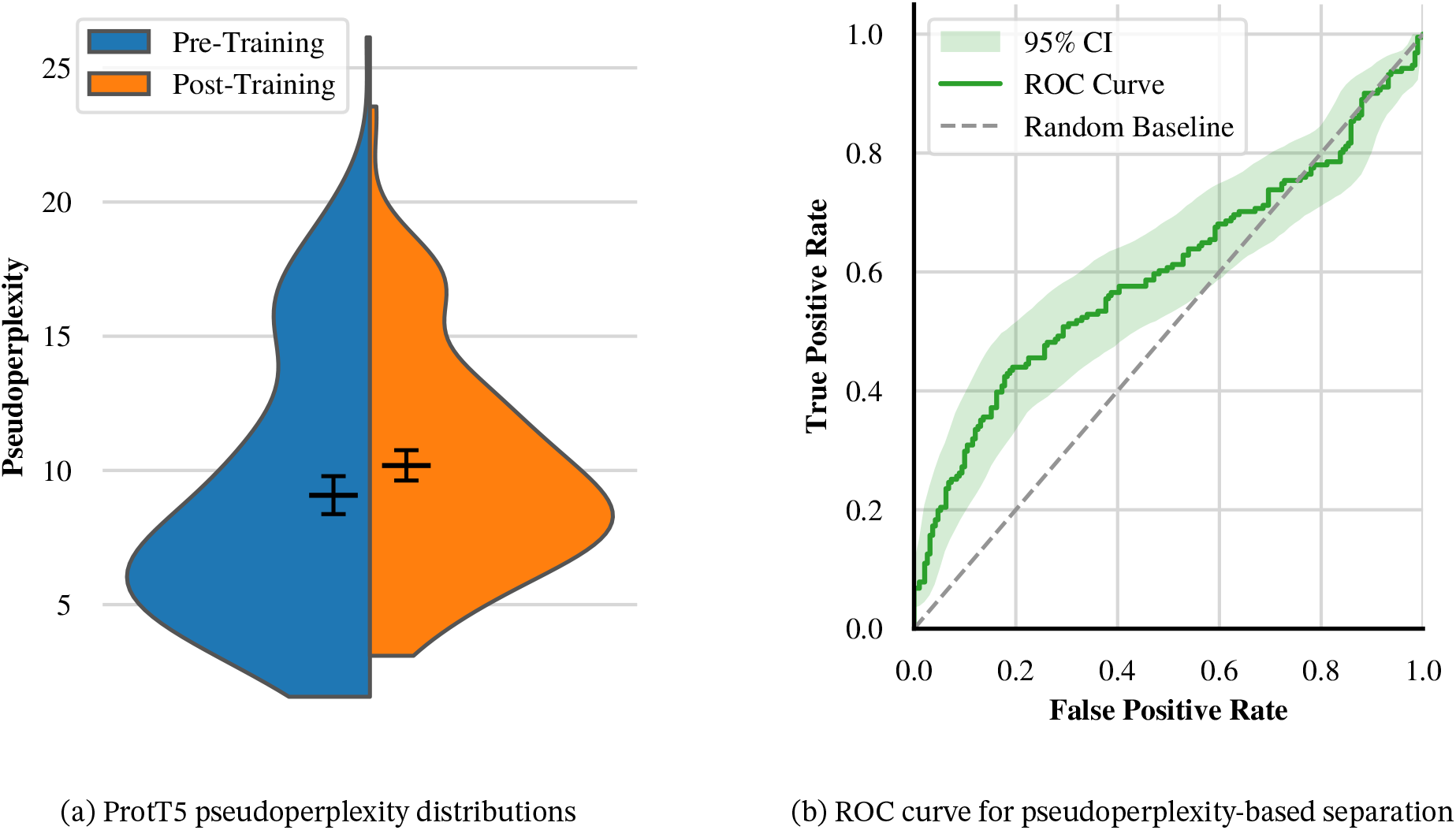
ProtT5 memorized a statistically significant yet modest portion of its training data. **(a)** The pseudoperplexity distributions for the matched pre-training and post-training datasets are shown as a split violin plot, with each half displaying the bootstrapped mean and 95% confidence interval (CI). The bootstrap mean for the pre-training dataset was 9.07 (CI95: (8.37, 9.79)), that for the post-training dataset was 10.18 (CI95: (9.62, 10.75)). Both distributions appeared bimodal, with most sequences concentrated in the lower and fewer in the upper mode. The upper mode was predominantly driven by sequences shorter than 50 amino acids. **(b)** The ROC curve was obtained by using pseudoperplexity as a binary classifier to separate pre-from post-training sequences. The dashed grey line indicates the random baseline corresponding to an AUC of 0.5. The calculated AUC of 0.6 with a 95% confidence interval of (0.54, 0.66) lies clearly above this baseline, indicating a detectable but modest training-data preference.

To quantify the extent of memorization, we performed an AUC-ROC analysis using pseudoperplexity as a binary classifier (see Section 3.4). The AUC of 0.60 (CI95: (0.54, 0.66), Figure 1b) obtained for ProtT5 fell above the random baseline of 0.5. For about 60% of the randomly drawn pre- and post-training protein pairs, the pseudoperplexity was lower for the pre-training protein. This confirmed that ProtT5 assigns systematically different likelihoods to trained-on and novel sequences. However, the AUC was still much closer to the random baseline than to perfect separation.

A model with strong memorization would be expected to separate the two datasets much more clearly, with an AUC closer to 1.0. Therefore, the observed AUC indicates a detectable but modest memorization signal.

To further investigate whether ProtT5 treats pre- and post-training sequences differently, we calculated per residue embeddings for both datasets, obtain per-protein embeddings through pooling (averaging over all residues in a protein), and projected onto two dimensions using t-SNE^1^. The per-protein (pooled) embeddings of pre- and post-training datasets were largely mixed in embedding space without any clear separation (Figure 2b). In fact, the mix of the actual data did not visually differ from a random mix (Figure 2a). This suggested ProtT5 to treat the two datasets essentially alike. Proteins with higher pseudoperplexity formed a distinct cluster in the upper right region of the embedding space (Figure 2c), suggesting that uncertainty is already encoded in the ProtT5 embeddings. This observation aligns with Prabakaran and Bromberg^1^, who reported that, in ProtT5, uncertain or underlearned protein embeddings can occupy latent-space regions that overlap with embeddings of randomly shuffled, non-biological sequences.

**Figure 2:**
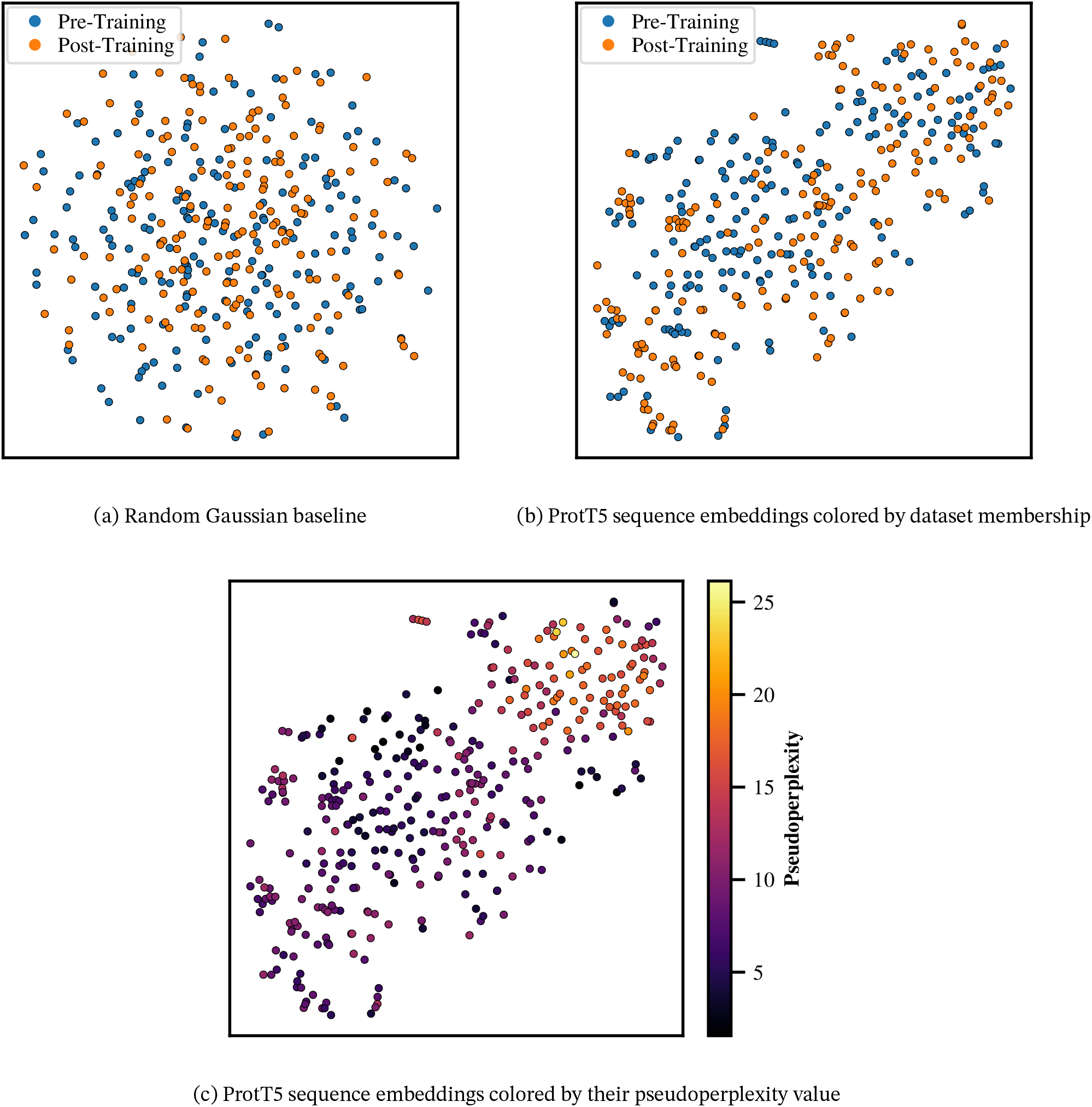
Pre- and post-training sequences overlap in embedding space, and uncertainty is encoded in the embeddings. All projections used a t-SNE perplexity setting of 30. Note that all panels used the same projection, i.e., the same point is at the same 2D-position. **(a)** A random zero-mean, unit-variance Gaussian baseline is shown for reference. **(b)** The ProtT5 embeddings for pre- and post-training occupy largely the same region in the t-SNE projection, i.e., no clear boundary separates the two. This suggests that ProtT5 does not treat the two datasets fundamentally differently in embedding space. **(c)** The same embeddings as in panels **b** and **c** are now colored by pseudoperplexity value. Sequences with higher pseudoperplexity form a distinct cluster in the upper right region, showing that uncertainty is encoded in the embeddings.

### 4.2. Higher-order *n*-gram divergence confirmed post-training sequences are truly novel

To establish the novelty of the post-training dataset at the local sequence level, we evaluated the pre- and post-training datasets across a series of complementary metrics. We first evaluated *n*-gram models with orders ranging from *n* = 1 to *n* = 7 by computing perplexity on both the post-training and the distribution-matched pre-training dataset. The distribution-matched pre-training data constitute subsets of the *n*-gram training data (Section 3.2.1).

Figure 3 shows the resulting perplexity distributions across all model orders and both dataset splits.

**Figure 3:**
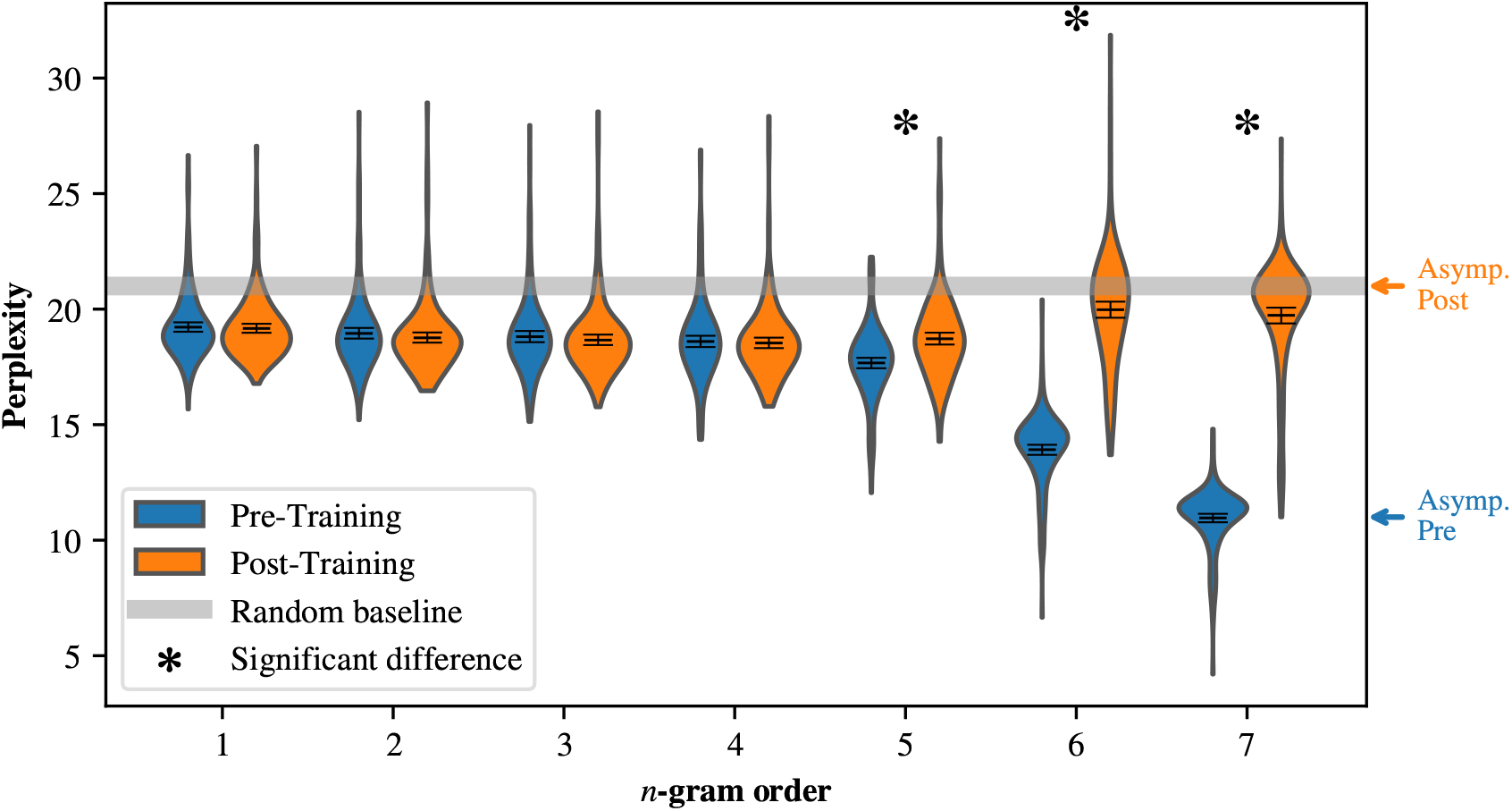
Pre-training perplexity dropped sharply at *n* ≥ 5 while post-training perplexity rose significantly at *n* = 6, marking the onset of overfitting to the training corpus. Perplexity distributions for *n*-gram models of order *n* = 1 to *n* = 7, evaluated on the post-training and distribution-matched pre-training dataset. The gray band at value 21 indicates the perplexity of a random baseline model, which intuitively chooses a token at random from a vocabulary of size 21. Arrows denote the theoretical asymptotic perplexity for pre- (= 11, blue) and post-training sequences (= 21, orange), as derived in the proofs in Supplementary 8.2 and Supplementary 8.3. Bootstrap means and 95% confidence intervals are shown as horizontal lines within each violin. Asterisks (*) denote statistically significant differences between the two datasets at the corresponding *n*-gram order after Bonferroni correction for seven comparisons.

For *n*-gram orders *n* = 1, …, 4, bootstrapped mean perplexity values ranged from 18.5 to 19.2 for both dataset splits (Figure 3, Table S2). By construction, these *n*-gram models correspond to maximum-likelihood estimates of conditional token probabilities based on training-set counts. The fact that perplexity remained relatively high — in close proximity to a random baseline model (=21) despite maximizing the likelihood of the underlying training set — indicates the inherent difficulty of amino acid prediction from local sequence context alone. Even with perfect local pattern matching, simple count-based models offer little improvement over random guessing, highlighting the complexity of this task.

The post-training empirical perplexities for *n* = 1, …, 4 were slightly lower (statistically insignificant) than those of the pre-training dataset. This may appear counterintuitive, given the novelty of the post-training data and their absence from the *n*-gram training set. We attribute this observation to the matching we applied to the pre-training distribution to make it as similar as possible to the post-training one. This rigorous matching makes the lower-order *n*-gram distributions (*n* = 1, …, 4) of the post-training dataset better match the *n*-gram frequency distributions of the training corpus than those of the pre-training dataset. This suggests that satisfying these rigorous matching criteria inherently restricts the pre-training dataset to a highly specific sequence subspace, making its local *n*-gram frequencies slightly less representative of the broader training dataset. Consequently, this non-significant difference in perplexity is an expected byproduct of our effort to render the two datasets as similar to each other as possible to eliminate potential confounders.

From *n* = 5 onward, the perplexity distributions of the two datasets differed significantly, and now the mean pre-training perplexity value was lower than that of the post-training (Figure 3, Table S2). The pre-training perplexities decreased, while the post-training perplexities did not differ significantly from those of the 4-gram model. The pre-post gap grew substantially at *n* = 6 and *n* = 7, where the mean pre-training perplexity dropped to 13.92 and 10.96, respectively, while the mean post-training perplexity rose, in a statistically significant way, to 19.98 and 19.73, indicating that the *n*-gram models overfit the training data beyond this point.

To further understand why the gap in perplexity became statistically significant exactly at *n* = 5, we analyzed theoretical *n*-gram space coverage, Jaccard similarities, and out-of-vocabulary (OOV) rates, each providing a unique perspective on the analysis.

First, to establish the coverage of our *n*-gram models, we examined the theoretical *n*-gram space and its empirical coverage. The space of possible *n*-grams grows exponentially with order *n*, bounded by Θ(|Σ|^*n*^), where |Σ| = 20 represents the amino acid tokens in our vocabulary (derived in Supplementary 8.4). The *n*-gram training set of approximately 68 million amino acid tokens covers > 99.4% of the *n*-gram space up to *n* = 4 (Table S1). However, at *n* = 5, coverage drops to roughly 91.5%, before declining sharply to ≈ 39.8% and ≈ 3.9% for 6-grams and 7-grams, respectively. This sparsity is severely amplified in the much smaller pre- and post-training evaluation datasets, due to their inherently limited size and diversity. Their coverage of the theoretical space drops from ≈ 20%(pre) and ≈ 22% (post) at *n* = 4 to merely ≈ 1.2%(pre) and ≈ 1.4%(post) at *n* = 5. However, this sparsity alone does not fully explain the divergence in perplexity between the two datasets.

Second, to establish that the new (post-training) proteins differed in their sequence from the old (pre-training), we calculated the Jaccard similarity — defined as the size of the intersection divided by the size of the union — between the distinct *n*-grams of pre- and post-training datasets (Figure S6a). For unigrams, the Jaccard similarity was 1.0, which is expected and indicates that both datasets share the same underlying amino acids. The Jaccard similarity remained high for *n* = 2 and *n* = 3 (Figure S6a). This indicates that proteins from both datasets share roughly the same building blocks, composed of 2- and 3-amino acid segments. However, this does not necessarily indicate that they use these building blocks with the same frequency. At *n* = 4, the Jaccard similarity dropped to 0.21, which indicates that roughly 79% of the 4-grams were dataset-specific. Nevertheless, nearly all post-training 4-grams existed in the *n*-gram training corpus; only three 4-grams were missing (Figure S6b). The mean perplexity did not differ significantly between 3-grams and 4-grams. This suggests that the 4-gram model did not outperform the estimates of the 3-gram despite the additional context. This is unexpected because longer contexts typically capture more discriminative patterns. The lack of improvement indicates that, for both datasets, the frequency distributions of 3-grams and 4-grams are similarly aligned with the training corpus, and the added context does not yield a meaningful generalization.

This dynamic shifted entirely at *n* = 5. The Jaccard similarity collapsed to 0.018, meaning that the pre- and post-training datasets shared only ≈ 1.8% of their distinct 5-grams. The perplexity values between 4- to 5-gram did not differ significantly for post-training but they did for pre-training sequences (Figure 3). This decrease in pre-training perplexity rendered the difference statistically significant at *n* = 5. The out-of-vocabulary (OOV) rate in Figure S6b, defined as the fraction of *n*-gram tokens in the post-training dataset that are absent from the *n*-gram training dataset, was 0.48% for *n* = 5. The 5-grams used in the post-training were still so general that nearly all of these appeared in the *n*-gram training corpus. Therefore, the gap did not result from encountering unknown *n*-grams, but from a frequency mismatch. The frequency distribution of the 5-gram model resembled more that of the pre-training than the post-training dataset, resulting in significantly lower perplexity for the pre-than for the post-training set.

The point of overfitting was reached at *n* = 6 and continued at *n* = 7, where Jaccard similarities of 0.002 and 0.001 indicated that fewer than 1% of the distinct *n*-grams were shared between pre- and post-training. The post-training OOV rate increased sharply from 0.48% at *n* = 5 to 24.5% at *n* = 6, and then to 77.6% at *n* = 7. Starting with *n* = 6, the *n*-gram building blocks making up the proteins in the post-training dataset became increasingly dataset-specific. These *n*-grams were penalized most heavily by the Laplace smoothing term, driving the perplexity of higher-order *n*-grams toward the random baseline of 21 (Supplementary 8.3). Because the model encountered extremely rare or completely unseen higher-order *n*-grams, the model effectively reverted to uniform prediction over the vocabulary. The pre-training dataset, by contrast, was a subset of the overall *n*-gram training corpus and therefore contained no OOV *n*-grams. As *n* increased, and contexts became more unique, the model’s frequency estimates became increasingly accurate for the pre-training dataset, driving the perplexity of those sequences towards the theoretical asymptote of 11 (Supplementary 8.2).

Finally, the *n*-gram perplexity values reported here are not directly comparable to the ProtT5 pseudoperplexity values. The two quantities are computed under fundamentally different objectives and probability models. *N*-gram perplexity conditions only on preceding tokens, whereas pseudoperplexity conditions on the full bidirectional context, including subsequent residues. The additional information lowers pseudoperplexity systematically over autoregressive perplexity for any given sequence. The two metrics are, therefore, conceptually distinct and should be interpreted independently rather than compared directly.

## 5. Conclusion

We investigated the extent to which ProtT5 memorizes its training data by comparing pseudoperplexity between a pretraining proxy dataset and a temporal holdout of novel protein sequences, both derived from UniProt. After removing all sequences from the post-training holdout that had more than 30% pairwise percentage sequence identity to BFD and UniRef50 2019_01, we matched the pre-training and post-training sets in terms of sequence length, cluster size, and taxonomic family distributions, removing three potential confounders from the comparison.

The results indicate that ProtT5 assigns systematically higher likelihoods to sequences drawn from the training distribution than to novel sequences, as evidenced by a statistically significant difference in pseudoperplexity between the two datasets. An AUC-ROC analysis further quantified this training-data preference as modest (AUC = 0.60, CI95: (0.54, 0.66)), suggesting that ProtT5 has not strongly overfit its training data. Complementary t-SNE projections of the per-protein embeddings showed that ProtT5 did not treat pre- and post-training sequences fundamentally differently in the embedding space. However, sequences from either set with high pseudoperplexity clustered distinctly. Thus, uncertainty seemed already encoded in the ProtT5 embeddings.

Complementary analysis using *n*-gram language models assessed the degree to which the post-training sequences were novel. We demonstrated that the pre- and post-training datasets differed significantly in their higher-order *n*-gram distributions, both in their distinct *n*-gram composition and their frequencies. With increasing *n*, the post-training proteins became increasingly disjoint from the underlying *n*-gram training corpus. This confirmed that the post-training dataset was distinct from the pre-training data, and also from the broader *n*-gram training superset. These unseen “protein building blocks” again evidenced the novelty of the post-training sequences.

Several limitations should be noted. First, our analysis is based on 191 sequences per dataset, which limits statistical power and the generalizability of effect size estimates. Second, we evaluated only ProtT5; whether these findings extend to other pLMs such as ESM-2 or autoregressive protein models remains to be determined. Third, our pseudoperplexity-based probe measures a statistical tendency rather than verbatim memorization in the sense of Carlini et al.16, and more targeted extraction-based approaches may reveal different memorization characteristics. Fourth, while the 30% sequence identity threshold used to exclude similar sequences from the post-training dataset is standard, remote homology below this threshold could still permit indirect information leakage. Finally, Feldman^1^ argues that some degree of memorization may be necessary for generalization on long-tailed distributions — a property that characterizes protein family abundances. The modest training-data preference we observe may therefore partly reflect necessary learning of rare but biologically valid sequence patterns rather than undesirable overfitting.

## 6. Author statements

## 6.1. Acknowledgements

We would like to thank Sarthak Parikh (LMU) for his ideas and support throughout the initial phases of this project. Thanks to Nikita Kugut, Dino Milijanic and Lothar Richter (all TUM) for help with administrative and hardware-related tasks. Last but not least, thanks to all who contribute toward making databases and programs publicly available. In particular to Alex Bateman (EBI Hinxton, England), Martin Steinegger (SNU Seoul, South Korea), and their teams.

## 6.2. Use of large language models

Parts of the code used in this work were developed with the assistance of large language models (Claude, ChatGPT, Gemini). LLMs were additionally used to assist in proofreading and improving the clarity of written text. All code and text were reviewed and verified by the authors.

## 6.3. Code and data availability

Code is released under the MIT License at https://github.com/aplaikner/prott5-memorization. Matched pre-/post-training sequences, ProtT5 embeddings, and pseudoperplexity values are available on the Hugging Face Hub under the `prott5-memorization` organization (https://huggingface.co/prott5-memorization).

The data and code are additionally hosted on Zenodo under https://doi.org/10.5281/zenodo.20591055 and https://doi.org/10.5281/zenodo.20594690, respectively.

## 8. Supplementary

### 8.1. Hardware

Pseudoperplexity computation and embedding calculation for both the pre- and post-training datasets were performed using the ProtT5-XL-U50 model. Both the pseudoperplexity calculation and the extraction of the per-protein sequence embeddings were executed on a single NVIDIA A100 GPU with 16 bit floating point precision.

As for the *n*-gram models, training and evaluation were conducted on the CPU of an Apple M1 MacBook Pro (8 core CPU, up to 3.2 GHz clock speed) with 16 GiB of unified RAM, using 64 bit floating point precision.

### 8.2. Asymptotic perplexity for seen sequences

Assume a finite training dataset *A*, and let *B* ⊆ *A* be a validation set consisting of sequences observed during training.

Consider a Laplace-smoothed *n*-gram model. For a finite dataset *A*, as *n* → ∞, *n*-grams become increasingly specific. For any fixed position in a sequence *x* ∈ *B* ⊆ *A*, there exists a threshold *n*_0_ such that for all *n* > *n*_0_, the corresponding *n*-gram occurs exactly once in the dataset. Thus, for sufficiently large *n*:

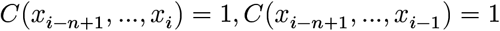

Applying Laplace smoothing:

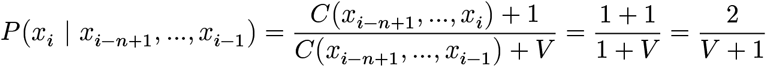

Thus, perplexity becomes:

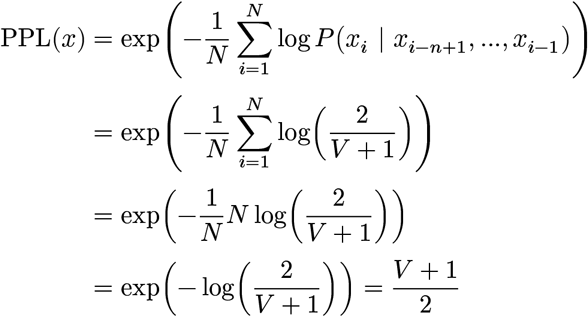

### 8.3. Asymptotic perplexity for unseen sequences

Assume a finite training dataset *A*, and let *C* be a validation set consisting of sequences not observed during training (*C* ∩ *A* = ∅).

Consider a Laplace-smoothed *n*-gram model. As *n* → ∞, contexts of a sequence *x* ∈ *C* become so specific that they do not appear in the training data. Thus, for sufficiently large *n*:

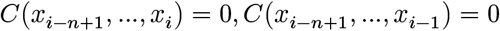

Applying Laplace smoothing:

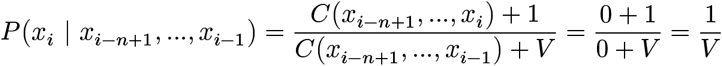

Thus, perplexity becomes:

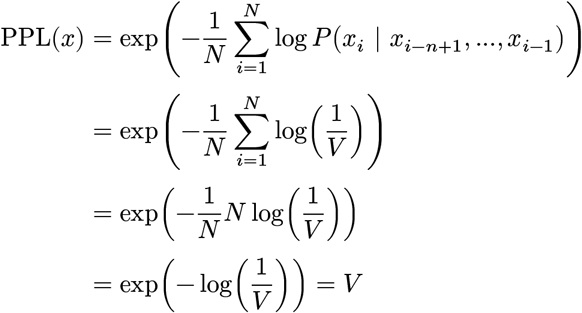

This corresponds to the perplexity a model would produce when sampling uniformly at random from the vocabulary.

### 8.4. Derivation of total *n*-gram space

We define the set of valid *n*-grams over an amino acid alphabet Σ of size |Σ| = 20 (standard amino acids), augmented with a start-of-sequence token <s> and an end-of-sequence token </s>. By construction, <s> may only appear as a contiguous prefix of the *n*-gram, and </s> may only appear as the final token. A valid *n*-gram, therefore, begins with *j* copies of <s> (where 0 ≤ *j* ≤ *n* − 1), followed by zero or multiple amino acid tokens drawn from Σ, optionally terminated by a single </s>. For a given prefix length *j*, the number of valid *n*-grams is 20^*n*−*j*^ (without </s>) plus 20^*n*−*j*−1^ (with </s>), giving 21 ⋅ 20^*n*−*j*−1^ combinations. Summing over all prefix lengths:

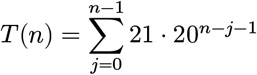

We introduce the index *m* = *n* − *j* − 1. As *j* runs from 0 to *n* − 1, *m* runs from *n* − 1 down to 0. Because summation is commutative, we can rewrite the sum over *m* = 0 to *n* − 1:

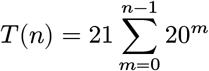

This is a finite geometric series. The formula of the geometric sum is:

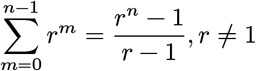

With *r* = 20, we obtain:

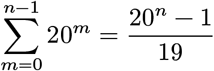

Thus, the final closed form for the number of possible *n*-grams is:

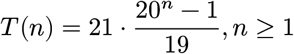

We can verify this formula for small *n*:

- 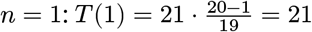 (standard amino acid tokens plus EOS token)
- 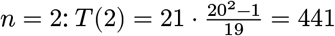

The *n*-gram space therefore grows exponentially as Θ(|Σ]^*n*^).

**Figure S1:**
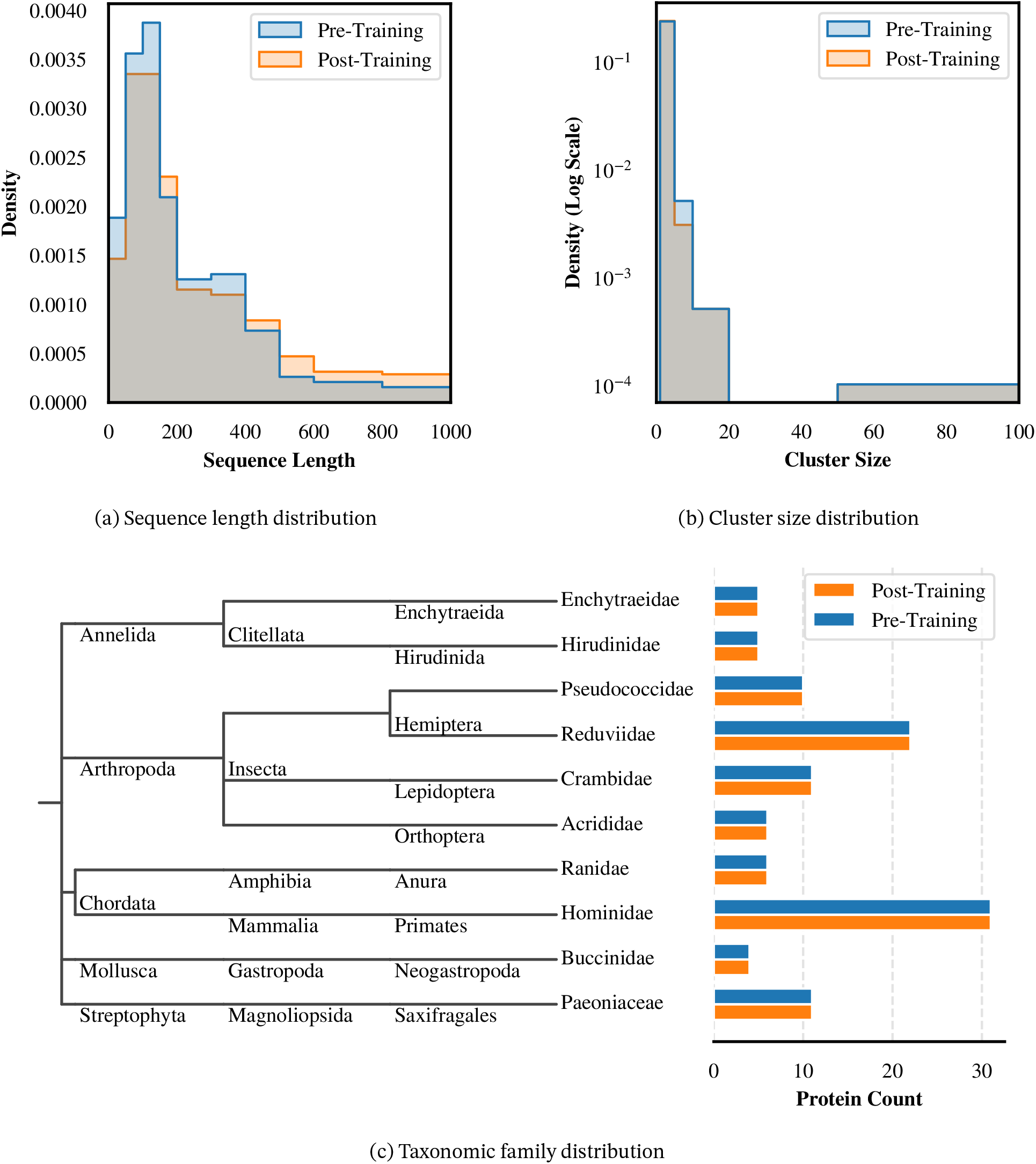
The matched pre-training and post-training datasets closely mirror each other across all three strata, confirming that the distribution matching procedure was effective. **(a)** The sequence length distributions are shown with the grey shaded area highlighting the overlap between the matched pre-training and post-training datasets. While small absolute discrepancies are present across all bins, the largest relative differences are observed in the (800, 1000] (6 pre vs. 11 post), (500, 600] (5 vs. 9), and (600, 800] (8 vs. 12) bins. **(b)** The UniRef50 cluster size distributions, shown on a logarithmic density scale, align almost perfectly, with only a single sequence difference in the (0, 1] and (1, 5] bin. **(c)** Taxonomic family distributions and hierarchical tree for the 10 most represented families in the post-training dataset, all of which are perfectly matched.

**Figure S2:**
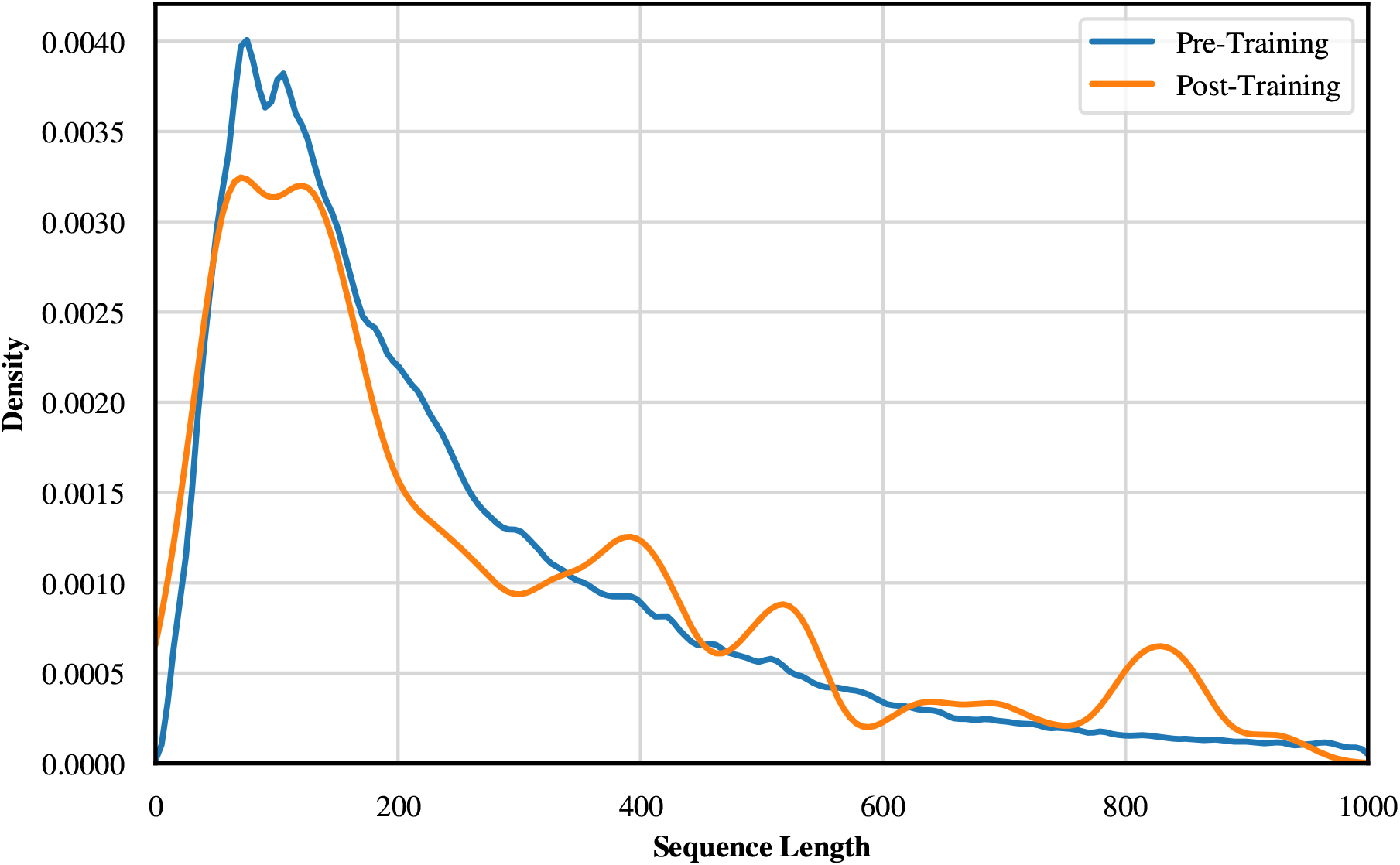
Most sequences fall between 50 and 200 amino acids, motivating finer binning in this range for distribution matching. Sequence length distribution of the filtered pre-training and post-training datasets. To reflect this concentration, the sequence length bins used for distribution matching were chosen to be narrower in the 50 to 200 amino acid range, with a bin width of 50 amino acids, and progressively broader above that threshold, with bin widths ranging from 100 to 200 amino acids. The upper bound of 1,000 amino acids follows directly from the length filter applied during dataset preparation (see Section 3.1.3).

**Figure S3:**
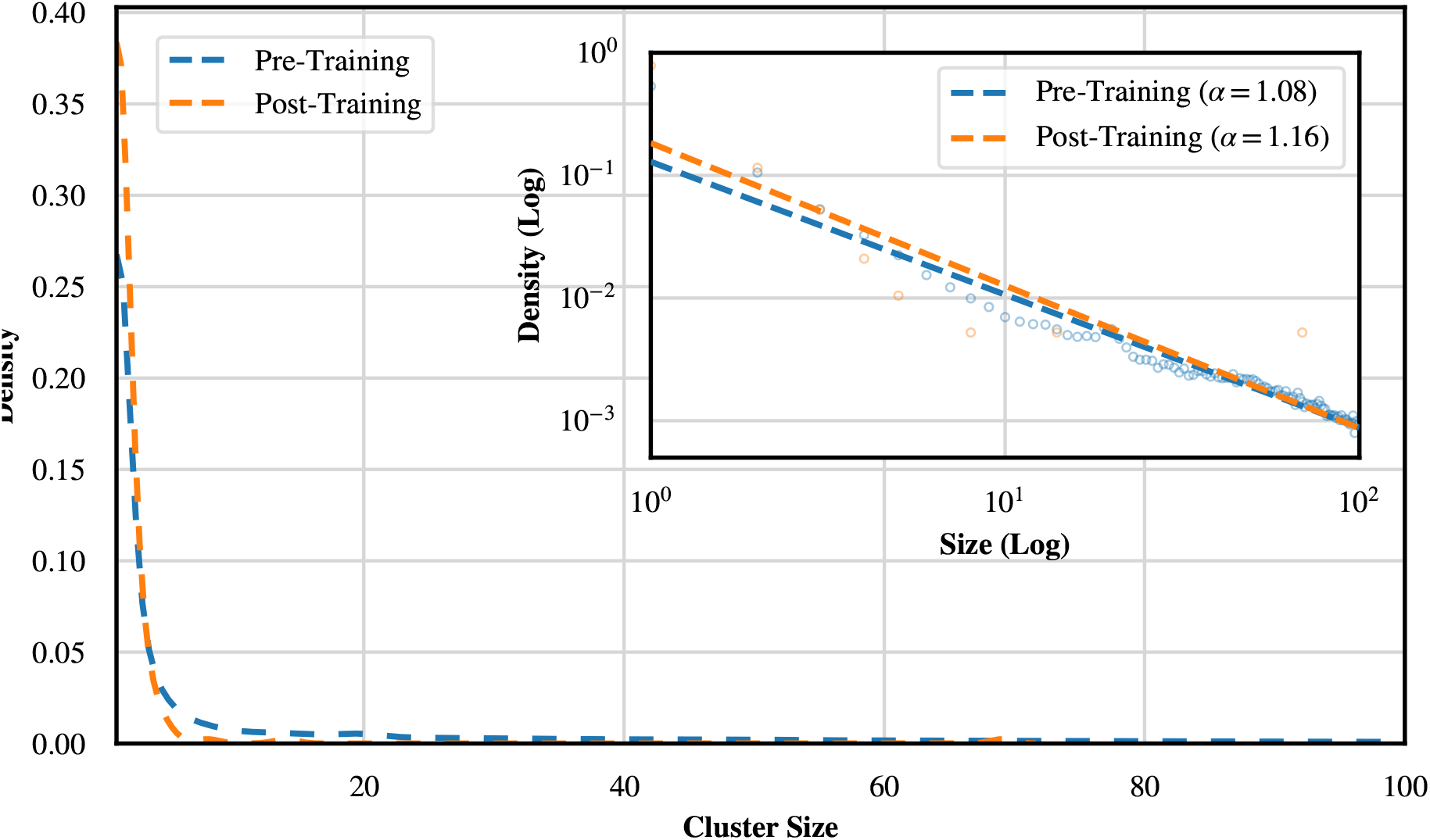
Cluster sizes follow a Zipf distribution concentrated at size 1, motivating finer bins in the lower range. UniRef50 cluster size distribution of the filtered pre-training and post-training datasets. The upper bound was set to 100, as the largest cluster size observed in the post-training dataset is 69, with a slightly higher threshold chosen to also accommodate pre-training sequences with larger cluster sizes. The inset displays the same data on a log-log scale as individual scatter points, with dashed lines showing the fitted Zipf curve. A Zipf distribution is a distribution where the frequency of an element is inversely proportional to its rank, often observed in biological data. The estimated parameter *α* characterizes the power-law decay of the cluster size distribution according to the relation *P* (*x*) ∝ *x*^−*α*^.

**Figure S4:**
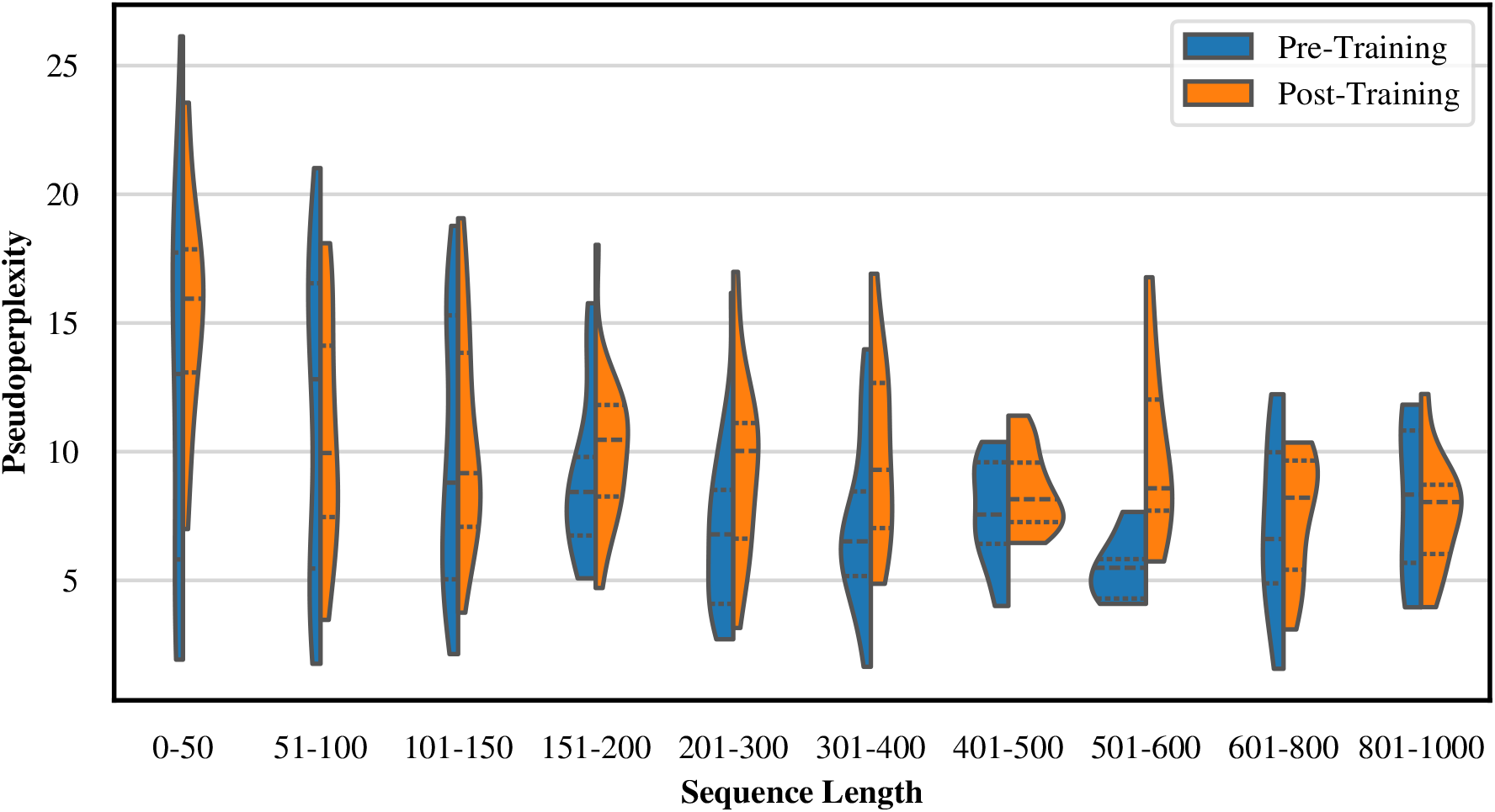
Short sequences under 50 amino acids are the primary driver of the bimodal pseudoperplexity distribution. Pseudoperplexity distributions of the matched pre-training and post-training datasets, grouped by sequence length bin, showing the median as well as the first and third quartile. Note that the per-bin distributions are based on very few sequences and should therefore be interpreted with caution. For all bins covering lengths from 151 to 1000 amino acids, the median, Q1, and Q3 of both datasets consistently fall in the lower pseudoperplexity mode observed in Figure 1a. The bin covering sequences of length 0 to 50 amino acids shows a different behavior, with a substantial portion of data points lying in the upper mode for both datasets, suggesting that very short sequences are the primary driver of the secondary mode. The bins of 51 to 100 and 101 to 150 amino acids mark a gradual transition, with data points falling inside both modes.

**Figure S5:**
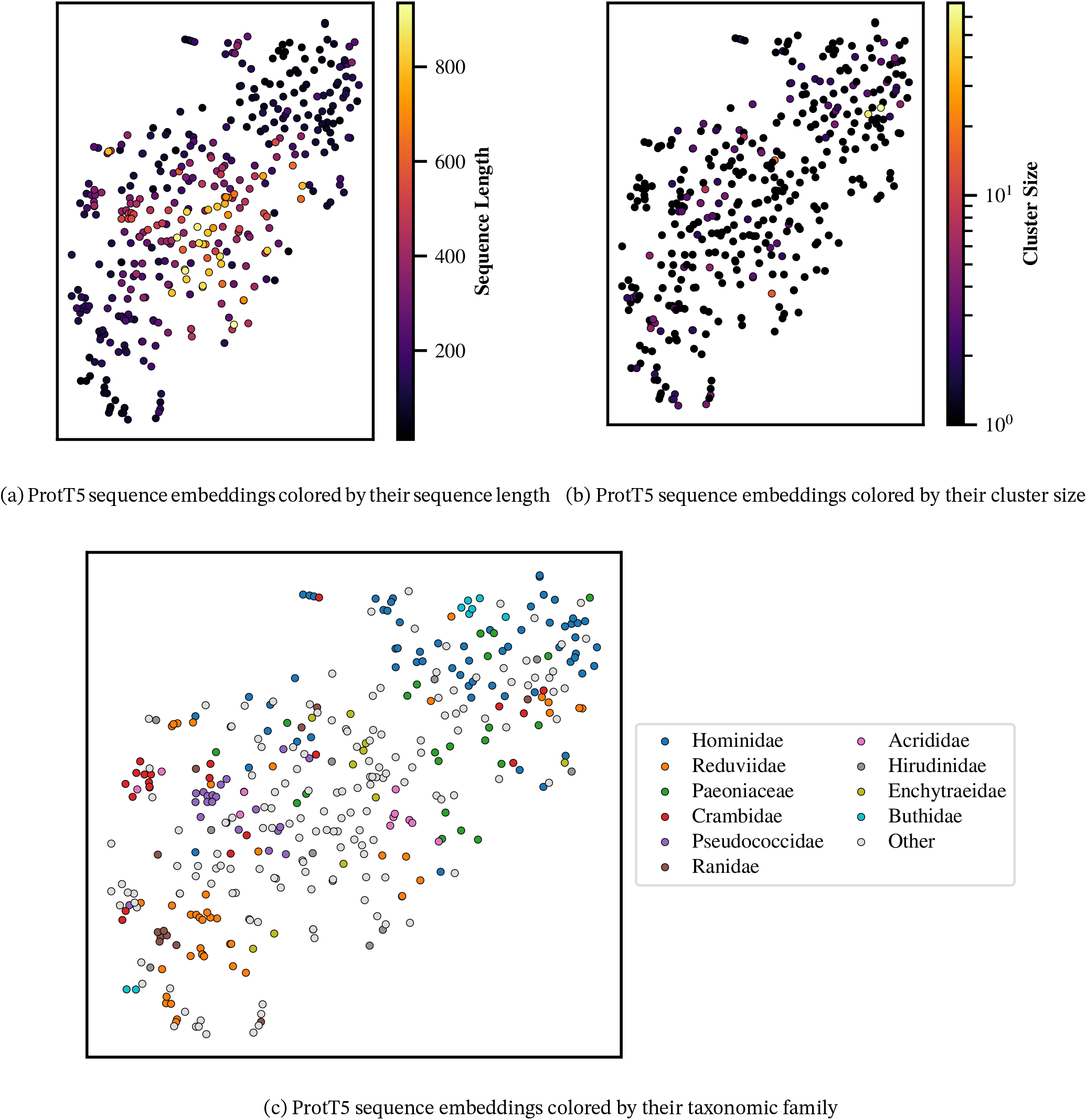
Sequence length is partly reflected in the ProtT5 embeddings, while cluster size and taxonomic family show no clear global structure. All panels show the same t-SNE projection of the matched pre- and post-training ProtT5 embeddings, using a t-SNE perplexity setting of 30. Thus, the same point has the same two-dimensional position in all panels. **(a)** Coloring by sequence length shows that longer sequences tend to concentrate in the central region of the embedding space, whereas shorter sequences are more broadly distributed. **(b)** Coloring by cluster size shows no clear spatial structure. Most sequences have small cluster sizes, and larger clusters do not form a distinct region in the projection. **(c)** Coloring by taxonomic family shows local clusters for some families. However, these clusters do not contain all sequences of the respective families. Thus, the projection does not show clearly separated, self-contained family-specific clusters, but rather small family-specific groups distributed across the embedding space.

**Figure S6:**
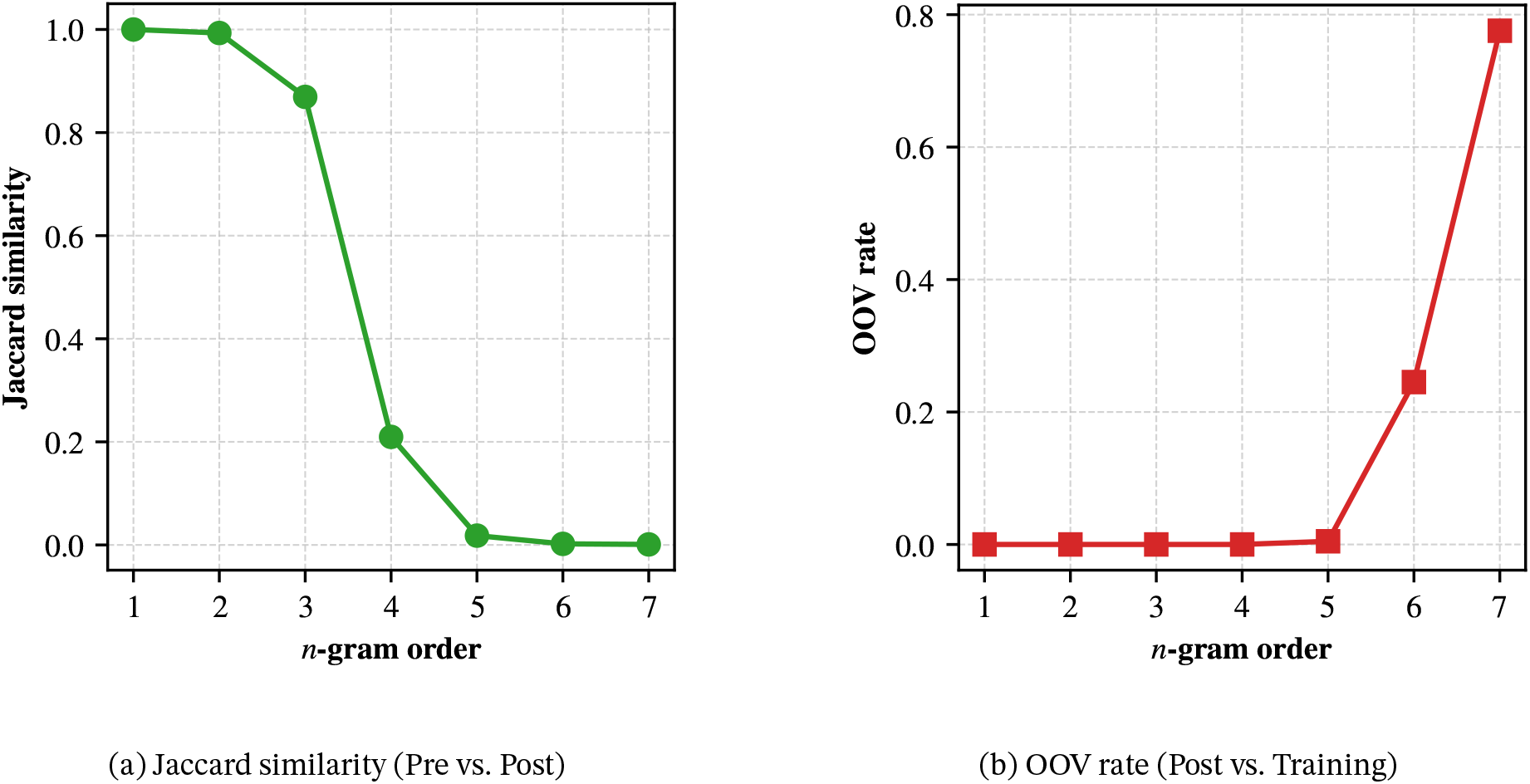
Post-training sequences occupy increasingly disjoint *n*-gram space at higher orders, confirming local sequence-level novelty. **(a)** Jaccard similarity of distinct *n*-grams between the post-training dataset and the distribution-matched pre-training dataset, and **(b)** *n*-gram out-of-vocabulary (OOV) rate of the post-training dataset relative to the *n*-gram training dataset, across *n* = 1 to 7. Jaccard similarity measures the overlap in unique *n*-grams, while OOV rate measures the fraction of *n*-gram tokens in the post-training dataset that are not observed in the training dataset. The sharp drop in Jaccard similarity at *n* = 4–5 indicates that the two datasets occupy largely disjoint regions of *n*-gram space. This is accompanied by a strong increase in OOV rate at higher orders, showing that longer sequence patterns in the post-training dataset increasingly fall outside the *n*-gram training distribution, consistent with sequence-level novelty.

**Table S1:** *N* -gram space coverage drops from near-complete at *n* ≤ 4 to below 4% at *n* = 7. Number of distinct *n*-grams observed in each dataset (*n*-gram training dataset vs. distribution-matched pre-training vs. post-training) and their coverage of the total possible *n*-gram space as derived in Supplementary 8.4. Note the decrease of coverage in all datasets as the *n*-gram order increases, reflecting the increase in *n*-gram sparsity.

| $n$ | Possible | Training | Pre-training | Post-training |
| --- | --- | --- | --- | --- |
| 1 | 21 | 21 (100.00%) | 21 (100.00%) | 21 (100.00%) |
| 2 | 441 | 440 (99.77%) | 439 (99.55%) | 438 (99.32%) |
| 3 | 8,841 | 8,820 (99.76%) | 7,634 (86.35%) | 7,671 (86.77%) |
| 4 | 176,841 | 175,889 (99.46%) | 35,282 (19.95%) | 38,928 (22.01%) |
| 5 | 3,536,841 | 3,235,860 (91.49%) | 43,726 (1.24%) | 50,029 (1.41%) |
| 6 | 70,736,841 | 28,128,370 (39.76%) | 44,535 (0.06%) | 51,269 (0.07%) |
| 7 | 1,414,736,841 | 55,555,742 (3.93%) | 44,680 (0.003%) | 51,545 (0.004%) |

**Table S2:** Pre- and post-training perplexities diverge significantly starting at *n* = 5, with the gap widening at higher orders. Bootstrap mean perplexity and 95% confidence intervals for *n*-gram models of order 1 to 7, evaluated on the post-training and distribution-matched pre-training datasets. Statistically significant differences between the two datasets (after Bonferroni correction) are marked with an asterisk.

| $n$ | Pre-training | Post-training | Significant Difference |
| --- | --- | --- | --- |
| 1 | 19.219 (19.022, 19.427) | 19.164 (18.976, 19.364) |  |
| 2 | 18.950 (18.725, 19.186) | 18.758 (18.550, 18.989) |  |
| 3 | 18.806 (18.570, 19.055) | 18.660 (18.445, 18.897) |  |
| 4 | 18.601 (18.359, 18.848) | 18.533 (18.313, 18.765) |  |
| 5 | 17.670 (17.443, 17.892) | 18.717 (18.467, 18.981) | * |
| 6 | 13.916 (13.692, 14.127) | 19.975 (19.627, 20.323) | * |
| 7 | 10.962 (10.774, 11.142) | 19.732 (19.381, 20.064) | * |

